# Cell motility influences microfluidics capturing in scRNA-seq

**DOI:** 10.1101/2025.02.21.639477

**Authors:** Lara López-Escobar, Ana M. Nascimento, Luisa M. Figueiredo

## Abstract

Microfluidic isolation methods for single-cell RNA sequencing (scRNA-seq) have primarily been designed for immotile cells, with limited consideration for motile cells, such as those with flagella or cilia. By studying the encapsulation efficiency of the flagellated *Trypanosoma brucei* using the 10x Genomics platform, we found that maintaining parasites at room temperature results in a low encapsulation yield. We implemented a rapid cooling method to 0ºC prior to encapsulation, which reduced parasite motility and preventing undesired transcriptomic change. This allowed for a representative scRNA-seq dataset, avoiding the disproportionate loss of the most motile forms. This study highlights the challenges of using motile cells in microfluidic systems and the biases caused by losing specific subpopulations, emphasizing the need for optimized protocols for non-standard mammalian cells.

**PLAIN LANGUAGE SUMMARY:** Single-cell RNA sequencing (scRNA-seq) is a powerful technique used to study individual cells, but most methods have been designed for cells that do not move. This can be a challenge when working with motile cells, like *Trypanosoma brucei*, a flagellated parasite responsible for sleeping sickness. In our study, we found that these parasites are poorly captured in microfluidic systems, such as the 10x Genomics platform, when kept at room temperature. Their movement reduces the efficiency of encapsulation, leading to biased results. To overcome this, we developed a simple cooling method that quickly lowers the temperature to 0°C before processing. This reduces parasite motility, improving capture rates while minimizing unwanted changes in gene expression. By applying this approach, we obtained a representative transcriptomic dataset, ensuring that all parasite forms were included. Our findings highlight the importance of adapting microfluidic techniques for motile cells to avoid losing key subpopulations and ensure accurate biological insights.

## INTRODUCTION

In the pursuit of unravelling cellular diversity, single-cell RNA sequencing (scRNA-seq) has emerged as a transformative tool. scRNAseq is defined as an advanced technology, used to measure the gene expression of individual cells. It comprises several steps, like cell isolation, library preparation, sequencing and data analysis. Studying single cell transcriptomes has enabled researchers to not only study gene expression in thousands of individual cells simultaneously^1^ but also to understand heterogeneity within diverse cell populations, such an antibiotic-associated cellular states under antibiotic perturbation in bacteria^2^ or a unique immune response against malaria parasite^3^.

A variety of methods have been developed for scRNAseq^4^. The central difference between them lies in how individual cells are isolated and barcoded. Microfluidics and droplet-based technologies eliminated the need for sorting and fixing cells, reducing cell stress during preparation, minimizing potential changes to gene expression and also greatly improving cell isolation efficiency^5^. In 10x Genomics (one of the most used technologies) microfluidics are used to encapsulate individual cells within nanoliter-sized droplets, along with Gel Beads and an oil emulsion, which are called GEMs. These Gel Beads carry two different barcodes: one called 10X, unique for each cell, and the other called Unique Molecular Identifier (UMI), unique for each transcript, allowing a more accurate quantification of transcript levels and enabling the elimination of artifacts such as PCR duplicates^6^.

Some cell characteristics pose challenges for encapsulation and barcoding. For example, yeast cells have a rigid cell wall that needs to be lysed with special enzymes^7^, and cells larger than 50 µm cannot pass through some microfluidic channels^8^. Since scRNAseq is relatively new (first paper was published in 2009^9^) the sample preparation has been reported already as a source of artifacts, and leading to false discoveries, related with stress-related genes expression^10^. The solution for this has been to keep the cells as inactive as possible, which is achieved by keeping them in cold before encapsulation^11^ or cryopreserve them^12^, as typically done bulkRNAseq. Another obstacle could be cell motility; however, this issue has not been widely reported, likely because cell loss is typically linked to washes or other technical problems^13^, making it difficult to demonstrate.

There are several motile cells in nature. Mammals have the highly motile sperm cells. Some immune cells also exhibit motility, though at much lower rates—such as naïve T cells at 6.2 µm/min, or CD8 T cells at around 4.3 to 5.2 µm/min^14^. On the other hand, among parasites, Leishmania^15^ and Trypanosoma^16^, for example, are highly motile. *Trypanosoma brucei*, a highly motile flagellated parasite^17^, crosses barriers in different hosts, including blood vessels in mammals^18^ and the proventriculus (or cardia) in the tsetse fly^19^. Highly motile behaviour is also detected in culture. The flagellum is essential for motility and plays a pivotal role in multiple facets of development, transmission, and pathogenesis^20,21^. The displacement of the flagellum also influences the parasite’s morphology and consequent speed of displacement^16,20^. In the mammalian host, quiescent stumpy forms have a shorter flagellum and slower velocity (10-20 µm/s) compared to proliferative slender forms, with a longer flagellum and higher velocity (20-50 µm/s)^22,23^. Additionally, parasites capable of reaching the brain and crossing the blood– cerebrospinal fluid barrier^18^ exhibit longer free flagella and increased velocity (50-100 µm/s)^23,24^. Similarly, in the tsetse fly, life cycle stages and flagellum length influence swimming velocity^23^.

In this study, we investigated how cell motility affects single-cell isolation and barcoding in a microfluidic system (10x Genomics). By examining two different stages of *Trypanosoma brucei* parasites with varying motilities, we observed that highly motile slender forms were underrepresented in the final dataset compared to the less motile stumpy forms. We found that immobilizing the parasites prior to encapsulation with rapidly cooling enhanced cell retention of motile parasites by up to 70% compared to when parasites were stored at room temperature. This approach results in a representative scRNA-seq data and underscores the importance of optimizing protocols for non-standard cells in microfluidic devices.

## RESULTS

### Parasites at room temperature are less efficiently encapsulated

To assess how *T. brucei* motility affects encapsulation in GEMs and vice versa, parasites were isolated from mice on day five post-infection (mostly slender forms^25^) and kept at room temperature (RT) until encapsulation. Encapsulated parasites were observed under an optical microscope, and we noticed that the physical process of encapsulation did not compromise the morphology of the parasites and individual motility (Video 1). However, some parasites showed a long displacement and high velocity, suggesting some parasites may have not been properly encapsulated or they erupted from the GEMs (Video 2).

To increase the probability of parasites being properly encapsulated, we tested whether rapidly lowering the temperature would decrease parasite motility. For that, a parasite suspension was rapidly cooled by submersion in a bath of 100% EtOH with dry ice for a few seconds, reaching 0 degrees Celsius, followed by immediate placement on ice, not letting the parasites freeze. This quenching is intended to rapidly reduce the metabolic activity and therefore the motility, while preventing cold-induced stress. This method is very commonly used in metabolomics to avoid changes in the metabolome of the cells ^26^. As a control, an equal amount of parasite suspension was left at room temperature (RT). As expected, we found that QUENCHED parasites showed a severe impairment in motility, with the parasites appearing almost immotile (Video 3) in contrast to parasites at RT, which showed a typical beating of the flagellum (Video 4).

To test if differences in motility would translate into differences in cell recovery, parasites were isolated from blood six days post-infection and split into two groups: QUENCHED or RT. For each condition, we prepared two technical replicates. The four samples underwent encapsulation, cDNA amplification and processed for library preparation: QUENCHED 1, QUENCHED 2, RT1, RT2 (Figure 1A). Libraries were sequenced by Illumina. Sequencing reads were aligned with CellRanger. After alignment, we already observed some differences in quality between QUENCHED and RT samples (Table 1). These numbers come directly from the raw matrix output from CellRanger, meaning that all barcoded cells are included. However, it is interesting that this raw data already shows differences, particularly in the number of genes per cell, which is twice as high in the QUENCHED samples than in RT samples (Table 1).

**Table 1.** Raw data vs filtered data. The table presents the number of genes per cell, unique molecular identifiers (UMIs) per cell, and total cell count for each sample before (RAW DATA) and after quality control (AFTER QC). “QUENCHED1” and “QUENCHED2” refer to samples processed using the quenching method, while “RT1” and “RT2” were prepared using a standard room-temperature (RT) approach. QC filtering significantly reduced the total number of cells while increasing the detected genes and UMIs per cell, indicating improved data quality.

| Sample | RAW DATA |  |  | After QC |  |  |
| --- | --- | --- | --- | --- | --- | --- |
|  | Gene/cell | UMI/cell | Total cells | Gene/cell | UMI/cell | Total cells |
| QUENCHED1 | 91,83 | 123,69 | 70380 | 979 | 1554 | 1324 |
| QUENCHED2 | 102,35 | 137,26 | 84261 | 998 | 1536,5 | 1383 |
| RT1 | 54,83 | 91,71 | 15396 | 959 | 1905 | 145 |
| RT2 | 59,23 | 99,76 | 84732 | 998 | 1612,5 | 698 |

**Figure 1.**
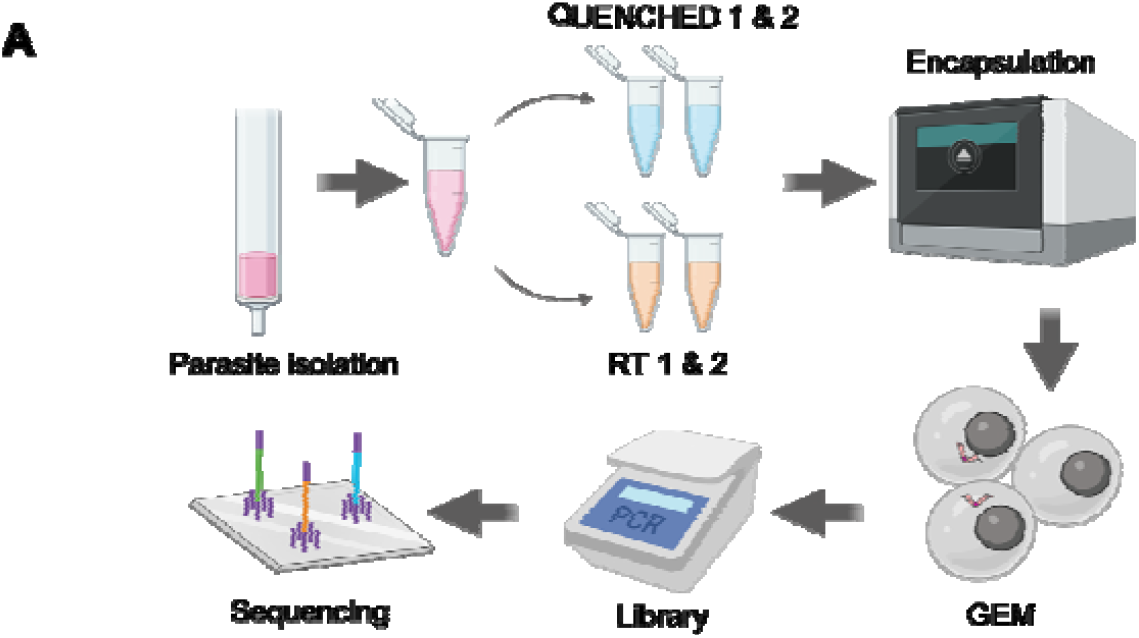
Motile parasites are less efficiently encapsulated. Parasites were isolated from the blood of three mice six days post-infection and purified through chromatography. Half of the parasite suspension underwent fast cooling to 0 degrees, using dry ice and ethanol (QUENCHED), while the other half remained at room temperature (RT). Both sets of samples underwent encapsulation using the Chromium controller, followed by library preparation and sequencing as recommended by 10x Genomics.

For quality control, we eliminated cells with fewer than 800 genes. This threshold was chosen based on previous studies analyzing bloodstream forms in scRNA-seq data, in which cells typically present around 1000 genes per cell^27–29^. We applied the same cutoff for both conditions since all cells originated from the same sample as technical replicates (Table 1).

After filtering, we found that the number of cells in QUENCHED samples was approximately threefold higher (1324 cells in QUENCHED1 and 1383 in QUENCHED2) than in RT samples (145 in RT1 and 698 in RT2) (Table1). From the initial input of approximately 7000 cells per sample, we recovered around 20% of cells in the QUENCHED samples and less than 6% in the RT samples. Given the low number of replicates (n=2), it is expected that the difference in recovery rates is not statistically significant. Consistently, during library preparation, we noticed a higher yield of cDNA (around 50% more) in the QUENCHED samples compared to the RT samples (Table S1). This suggests a more efficient encapsulation of QUENCHED parasites than those kept at RT, resulting in a great number of correctly barcoded parasites.

The recovery rate in 10x Genomics workflows is highly variable. While 10x advertises a recovery rate of around 60%^30^, significant heterogeneity is observed across studies, largely influenced by various technical conditions^31^. For *Trypanosoma brucei*, other studies have reported a wide range of recovery rates, from 16% to 40%^27–29^. In our study, the recovery rate from the QUENCHED condition (20%) falls within this range, whereas the RT condition shows a notable drop to 6%. We conclude that samples stored at RT yielded fewer GEMs with an acceptable transcriptomic cell profile than samples quenched or placed in ice^27–29^, thereby decreasing the final yield of the scRNA-seq experiment.

### Quenching improves retention of slender forms

Given stumpy forms are less motile^22^, we wondered if stumpy forms are more efficiently encapsulated than slender forms. Given that a parasite population isolated from blood six days post infection is typically composed of a mixture of slender and stumpy forms^25^, we used the four samples from this study to compare the encapsulation yield of slender and stumpy forms in each sample.

To confirm that our four samples contained a mixture of slender and stumpy forms, we merged the four datasets and visualized them using Uniform Manifold Approximation and Projection (UMAP)^32^ reduction. The four merged samples revealed the presence of two clusters (Figure 2A), which correspond to slender and stumpy forms using previously established gene signatures for these two life cycle forms (Table S2)^27^ (Figure 2B). Plotting the expression of three of these marker genes (PAD2, ZC3H20 and PYK1) help us to visualise the two life cycle stages (Figure 2C). We noticed that the actively transcribed Variant Surface Glycoprotein gene AnTat1.1E (manually added to the reference genome along with other VSG genes) was downregulated (Figure 2C, Figure S1A), as previously reported^33^ in bulk RNAseq data. Thus, this gene can also help the validation of slender/stumpy stages early in infection when VSG switchers are not detected in scRNAseq data. We also observed differences in UMI and gene counts between the two stages, with the stumpy forms showing a higher number of detected genes and transcripts than the slender forms (Figure S1B). Although this may seem counterintuitive, the downregulation of the active VSG in stumpy forms may desaturate the sequencing, allowing for the detection of a higher number of genes in stumpy forms that in slender forms.

**Figure 2.**
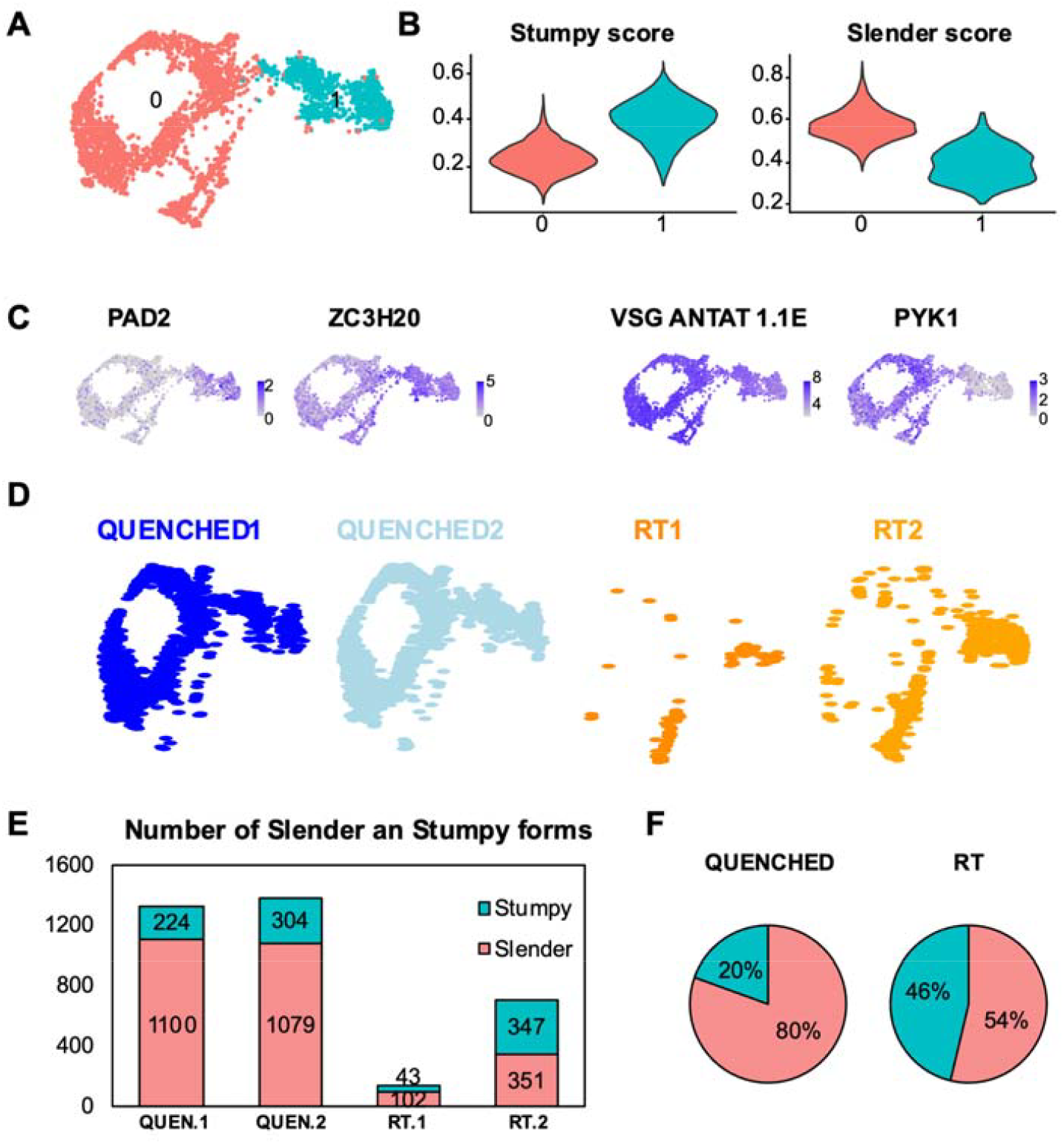
Quenching improves retention of slender form parasites. **(A)** Uniform Manifold Approximation and Projection (UMAP) plot of four merged samples showing two unsupervised clusters (cluster 0 in red and cluster 1 in cyan). **(B)** Violin plots from panel (A) that show the score of the cells from cluster 0 or cluster 1 as “stumpy” or “slender” with the genes showed in Tabel S1. The “slender score” and “stumpy score” consist of the average expression of associated marker genes for each cluster identified in A. **(C)** UMAPs coloured by transcript counts for two stumpy markers (EP1 and PAD2) and two slender markers (PYK1 and VSG AnTat1.1E). Scales show raw transcript count per cell. **(D)** Same UMAP plot from panel A but divided in the four idents that composes it: in blue the QUENCHED samples and in orange the RT samples. **(E)** Number of parasites belonging to the stumpy (cyan) or slender (red) stages in the four samples QUENCHED1, QUENCHED2, RT1, RT2 (colored as panel A). **(F)** Pie chart indicating the percentage of slender and stumpy forms found in the sum of two replicates at each temperature condition.

Segregation of the UMAP based on the type of sample shows that RT samples have almost no cells in the slender cluster relative to QUENCHED samples (Figure 2D). In both replicates of the QUENCHED samples, we detected 224 and 304 stumpy forms and 1100 and 1079 slender forms (Figure 2E). Conversely, when parasites were incubated at RT, in the two replicates we detected 43 and 347 stumpy forms and only 102 and 351 slender forms (Figure 2E). This means that when parasites were kept at RT, the proportion of sequenced slender vs stumpy was 54:46%, which is markedly different from the 80:20% distribution observed in the quenched samples (Figure 2F). In other words, at RT we lost on average 79% of slender forms and 26% of stumpy forms relative to fast cooling conditions (453 vs 2179 total slender forms; 390 vs 528 total stumpy forms). Given that slender forms are more motile than stumpy forms^22^and motility is reduced at lower temperatures (Videos 3-4), quenching probably facilitated the recovery of less motile slender forms within the GEMs and thus the final sequenced population of parasites is more representative of the original population in the blood.

Overall, these results show that when parasites are stored at room temperature before encapsulation, slender forms are disproportionately lost from final encapsulated GEMs. This biased loss of a subpopulation is likely attributed to the higher motility of slender forms relative to stumpy forms^22,23^. Loss of motile forms can be avoided by immobilizing them using fast cooling before encapsulation.

### Slender forms are more sensitive to temperature changes than stumpy forms

Next, we evaluated the impact of the temperature on the transcriptome of slender and stumpy forms separately. To do so, we conducted a pseudobulk analysis of slender or stumpy form parasites in each temperature condition, followed by a correlation analysis of the transcript levels (measured as mean gene counts normalized with the total number of cells in each condition). We observed that the transcriptomes of slender forms in QUENCHED vs RT are highly correlated (correlation coefficients of 0.93). The transcriptomes of stumpy forms are also highly correlated 0.95 (Figure S2A and B). As expected, these correlations are both higher than the correlation between the transcriptomes of slender versus stumpy forms (0.91) (Figure S2C). These analyses indicate that fast cooling does not have a major impact on the transcriptome of neither slender nor stumpy forms.

However, we can detect differences between conditions, especially in the slender forms, where most RT parasites form an independent cluster (Figure 3A). To identify the genes affected by temperature in slender or stumpy forms, we performed a differential expression analysis and selected those with a log fold-change between conditions equal or higher than 0.5 and a p-value lower or equal than 0.05. In slender forms, we found 935 differentially expressed genes (from a total of 8930, i.e.∼ 10%) between QUENCHED and RT conditions, while in stumpy forms, 261 genes (∼3%) show differential expression (Figure 3A-B and Table S3 and S4).

**Figure 3.**
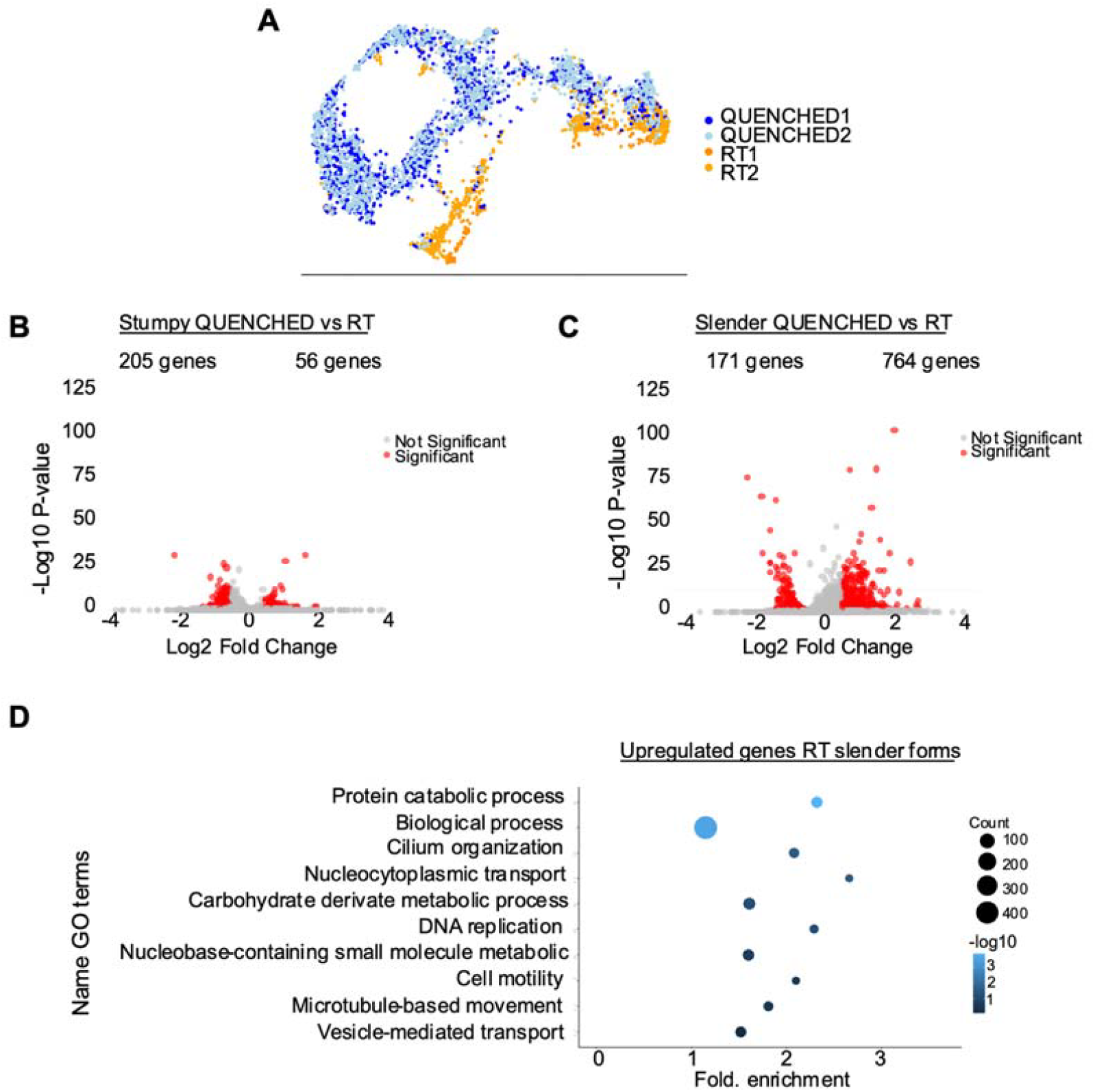
The transcriptome of slender forms is more sensitive to temperature variations than stumpy forms. **(A)** Uniform Manifold Approximation and Projection (UMAP) plot of four merged samples: in blue the QUENCHED samples and in orange the RT samples. **(B)** Volcano plot of differentially expressed (DE) genes of stumpy forms in QUENCHED versus RT conditions. Genes colored in red are significantly differentially expressed. **(C)** Volcano plots of differentially expressed (DE) genes of slender forms in QUENCHED versus RT conditions. Genes colored in red are significantly differentially expressed. **(D)** Gene ontology (GO) analysis of differentially expressed genes upregulated in slender form (panel B) of RT condition. The color of the dots represents the adjusted p value of each GO term: from dark blue (higher) to light blue(lower). The size of the dot represents the gene count in each GO term, ranging from the smallest dot (lowest count) to the largest dot (highest count).

Go term analysis of upregulated genes in slender forms, revealed an enrichment of terms involved in DNA replication (GO:0006260), nucleocytoplasmic transport (GO:0006913), and microtubule-based movement (GO:0007018), suggesting enhanced replication, protein transport, and cytoskeletal activity in RT conditions (Figure 3D). Importantly, no GO terms associated to stress were detected (such as GO: 0009409, GO:0070417). Amongst the downregulated genes in slender forms, we did not find a go term exclusive of this group of genes. This could be because of the low number of genes analysed. These findings confirm that RT conditions do not stop parasite proliferation and cellular activity, and thus parasites are more active than the ones quenched.

In summary, quickly dropping temperature does not have a major impact on transcriptome of slender forms, and even less on stumpy forms. Not surprisingly, in slender forms, quick cooling reduced the transcript levels of a few genes involved in parasite motility and proliferation.

## DISCUSSION

Over the past decade, single-cell RNA sequencing (scRNA-seq) methods have been significantly improved. Various single-cell isolation techniques, such as fluorescence-activated cell sorting (FACS)^34^, microfluidic droplets^5^, and emerging technologies like PARSE^35^ (which utilizes cell labelling and multi-pipetting) and BD Rhapsody^36^ (which employs microwells), have been developed. While these methods are effective, they are optimized for standard mammalian cells. In this paper we describe that quenching of parasites prior to microfluidics-based encapsulation (such as in 10x Genomics), immobilize parasites, helping with the cell recovery and avoiding major transcriptomic stress.

Protocols from 10x Genomics have been successfully been used in *T. brucei*^27,28,28,37^, often incorporating an ice incubation step prior to encapsulation, which makes the parasites more immotile than normal RT. However, significant loss of parasites—larger than typical mammalian cells—has been observed in several studies, with recovery rates ranging from 20% to 52% (22% in this study) while in mammalian cells recovery rates vary between 40% to 65%^30,38^. While other factors can be involved (size of the cell, for instance) we hypothesized that cell motility could be also involved.

Placing parasites on ice reduces their motility. However, because encapsulation happens at room temperature, parasites may partially recover their motility. In fact, previous studies using microfluidic have shown that *T. brucei* parasites survive and even divide for extended periods within droplets^39^. In this study we confirmed that encapsulating parasites at room temperature using the 10x Genomics microfluidic device does not induce changes in motility or morphology (Videos 1-2).

To reduce the chances that parasites would not recover their motility during encapsulation, while avoiding transcriptomic changes due to cold shock, in this work we decided to quench parasites (rapid cooling), a method traditionally used in metabolomics studies to reduce artifact changes associated to cold shock^40^. The only difference between quenching parasites or placing them on ice (standard protocol recommended by 10x Genomics), is the rapid cooling of the parasite to 0ºC on ethanol in a dry ice bath. Parasites are never frozen, and after 5-7 seconds on this bath, the tube is transferred to ice to keep parasites cold. While we did not compare quenching versus standard ice protocol side by side, by comparing our efficiency of cell recovery with published data, it seems the efficiencies are similar. Quenching has already been used in bulk RNA-seq in *T. brucei* to reduce cold shock^41^.

While analysing the transcriptomic data, we detected an interesting pattern that could be helpful to distinguish *T. brucei* life cycles stages with different levels of VSG expression. Indeed, we found that stumpy forms displayed a higher number of genes and transcripts per cell than slender forms. This is likely due to the differential expression of VSGs between these two life cycle stages. VSG mRNA is highly abundant in slender forms and thus many reads (7.5% of total reads in this dataset) align to this single gene, reducing the opportunity for other transcripts to be sequenced. When VSG expression is lower, other transcripts become more detectable, resulting in the detection of more genes/cell. This effect could explain why in other scRNA-seq studies of *T. brucei* tsetse fly stages, non-VSG-expressing stages such as procyclic forms and epimastigotes have a higher number of transcripts and genes per cell than VSG-expressing metacyclic forms (between 10%-50% higher)^28,37,42^. This finding could aid in the identifications of cell clusters in *T. brucei* scRNA-seq datasets.

While we did not find major transcriptomic differences between RT and QUENCHED conditions (Figure S2), 10% of genes were differentially expressed in the slender forms between the two temperature conditions (Figure 3A). Specifically, at RT we found upregulation of genes involved in motility and proliferation, and importantly we did not detect changes in gene expression that would suggest cells were under stress (Figure 3D). These results are consistent with the observations shown in the videos 1-4, in which the quenched parasites do not exhibit any morphological or behavioural changes.

While the data shows that when parasites were incubated at RT the capture efficiency was lower than when parasites were quenched, we cannot unequivocally attribute this to the encapsulation moment, as we were unable to observe the process continuously with a camera. However, we believe it is important to consider the implications of motility when working with highly motile organisms. Previous studies describing scRNA-seq on *Leishmania*^43^ or *Trypanosoma cruzi*^44^ have consistently included a step in which parasites were placed on ice prior to encapsulation, which seems to at least partially mitigate motility. These consideration should be taken into account in microfluidics-based analysis of other highly motile unicellular organisms, such as *Chlamydomonas reinhardtii*^45^, *Euglena gracilis*^46^, some Diatoms^47^ and bacteria as *Pseudomonas aeruginosa*^48^.

Exploring how scRNA-seq microfluidic devices selectively capture distinct cell populations within a heterogeneous sample is crucial for understanding the limitations and biases of these methods. Such investigations could lead to refining and optimizing scRNA-seq protocols, enabling more efficient accurate capture of cellular diversity across different biological contexts.

## MATERIALS AND METHODS

### Trypanosoma brucei cell lines

Mice were infected by *Trypanosoma brucei* EATRO1125 strain, AnTat1.1E clone, chimeric triple reporter cell-line ^49^expressing the red-shifted firefly luciferase protein PpyREH9, tagged with TdTomato and TY1. For in vitro scRNAseq analysis EATRO1125 AnTat1.1 GFP::PAD1utr was used, in which a GFP gene is followed by a PAD1 3⍰ UTR that confers maximum expression in parasites that have initiated differentiation to stumpy forms.

### Animal infections

Animal experiments were performed according to EU regulations and approved by the Órgão Responsável pelo Bem-estar Animal (ORBEA) of Instituto de Medicina Molecular and the competent authority Direcção Geral de Alimentação e Veterinária (licences 018889/2016 and 017549/2021). Mice were group-housed in filter-top cages in a Specific-Pathogen-Free barrier facility under standard laboratory conditions: 21 to 22°C with 45 to 65% humidity and a 12⍰h light/12⍰h dark cycle. Chow and water were available ad libitum. All infections were performed in 7–10-week-old wild-type male C57BL/6J mice with origin in Charles River Laboratories, by intraperitoneal (i.p.) injection of 2000 parasites.

### Parasite isolation from blood

After 5 or 6 days of infection (depends on the time point of the experiment), mice were sacrificed mice with CO_2_. Blood was removed by heart puncture (1 mL per mouse) and deposited in a 10 mL falcon with 2% CGA (Citrate Glucose Anticoagulant) and centrifuged for 10 min at 2000 rpm. The layer between the red blood cells and serum was removed by pipetting and mixed with 3 mL of the Separation Buffer from the DEAE sepharose™ Fast Flow column. After the selection with the DEAE column, parasites were counted using a disposable haemocytometer and diluted at the desired concentration in the buffer 1X PBS supplemented with 1% D-glucose (PSG) and 0.04% bovine serum albumin (BSA) at room temperature. For microscopy imaging, parasites were resuspended at 1-10 million parasites in 1 mL and for the scRNAseq experiment 1×10^5^ parasites were resuspended in 1 mL.

For the RT scRNA-seq protocol, parasites were left in a microtube with 1% D-glucose (PSG) and 0.04% bovine serum albumin (BSA) at room temperature until the moment of single cell isolation in the Chromium Controller (RT tubes). For the QUENCHED samples, 500 μL of the original tube at RT were transferred into another 1.5 mL microtube and immediately placed in a bath of 100% EtOH with dry ice for 5-7 seconds, until the thermometer marked 0 degrees. Next, this tube was transferred to a water bath with ice and kept there until the moment of single cell isolation.

### Single-cell RNA sequencing protocol

For each scRNA-seq sample, a total of 7000 cells from the mixed sample were loaded into the Chromium Controller (10x Genomics) to capture individual cells with unique barcoded beads. Libraries were prepared using the Chromium Single Cell 3⍰ GEM, Library & Gel Bead Kit v3.1 Dual index (10x Genomics). Total cDNA was measured with the LabChip GX Touch Nucleic Acid Analyzer and the DNA kit. Library preparation was performed following the protocol from 10X Genomics. Sequencing was performed with Novogene with the Illumina NovaSeq™ 500 platform to a total of at last 100Gb per sample. At least 20000 reads per cell was required for each sample.

### Single-cell RNA sequencing quality control

Illumina reads were aligned to the reference genome^27^ combined with the genes encoding for luciferase protein PpyREH9, TdTomato, TY1 tag^49^, GFP, the VSG AnTat1.1E (closest sequence GenBank: X01843.1) and a list of early expressed VSG ORF^50^. Reads were mapped onto the genome using cellranger-7.2.0 count function with the default settings. Low-quality cells were removed by filtering for low unique genes detected (<800), high proportion of kDNA (>2%) and high proportion of rRNA (>8%)^27^.

### Single-cell RNA sequencing normalization and clustering

The data was normalized following the procedure outlined in Briggs *et al*. 2021^27^. A low resolution (0.2) was selected for identification of the two most distinct populations. As the samples were sequenced together, no integration was performed to enable a comprehensive analysis of the differences between them. Replicates from the QUENCHED condition indicated the absence of batch effects in the data, with observed changes only attributable to biological factors.

### Stumpy and slender score

The list of genes from slender and stumpy form were generated from Briggs et al.2021^27^ data. This set of genes was utilized to calculate the average expression stumpy or slender scores using the AddModuleScore function from Seurat as indicated in Larcombe *et al*. 2023^29^.

### Pseudobulk analysis

All counts were aggregated to the sample level using “AggregateExpression” from Seurat, across four different samples: slender and stumpy from QUENCHED condition, and slender and stumpy from the RT condition.

### Microscope videos

To record parasite motility on video, 10 µl of parasites were placed on a slide. For the QUENCHED parasites, slides were pre-chilled on ice for at least 10 min before acquiring the videos rapidly, ensuring that the parasites remained cool and unaffected by laser temperature. To record parasite motility in GEMs, the training kit from 10XGenomics was utilized, and parasites were collected as described in the Parasite Isolation section of the bloodstream form. All the videos presented in this article were taken in a 3i Marianas SDC (spinning disc confocal) microscope (Intelligent Imaging Innovations, equipped with a Yokogawa CSU-X1 confocal scanner and a Photometrics Evolve 512 EMCCD camera)^51^ (https://imm.medicina.ulisboa.pt/facility/bioimaging/doku.php?id=3i_marianas_sdc). Transmitted light and laser line 561 nm were used to image the parasites and its TdTomato expression. The objectives used in these acquisitions were a 10x EC Plan-Neofluar (0.3 NA; 5.50 mm WD), and a 20x Plan-Apochromat (0.8 NA; 0.55 mm WD). A hundred images were obtained in each time lapse, with an acquisition rate of 2 frames per second. For all acquisitions, the software used was 3i Slidebook reader v.6.0.22 allowing export of images in TIFF format. TIFF documents were then processed using ImageJ 1.52a Java 1.8.0_112 [64-bit].

## FIGURE LEGENDS VIDEOSs

**Video 1. Parasite motility after GEM encapsulation (multiple focal planes)**. GEMs generated with 10XGenomics protocol are shown in blue (Phase contrast) and the parasites in yellow (TdTomato signal). It shows multiple layers of GEMs at 10X magnification at 500 ms. Parasites remain motile, and some of the GEMs move and burst.

**Video 2. Parasite motility after GEM encapsulation (single focal plane)**. GEMs are shown in grey (Phase contrast), and the parasites in red (TdTomato signal). This video shows a single layer of GEMs at 20X magnification at 500 ms speed.

**Video 3:** Reduced parasite motility after quenching to 0 degrees. Parasites express the fluorescent reporter, TdTomato, which is shown here in yellow.

**Video 4:** Regular parasite motility at room temperature (RT). Parasites express the fluorescent reporter, TdTomato, which is shown here in yellow.

## SUPPLEMENTARY MATERIALS

**Figure S1:**
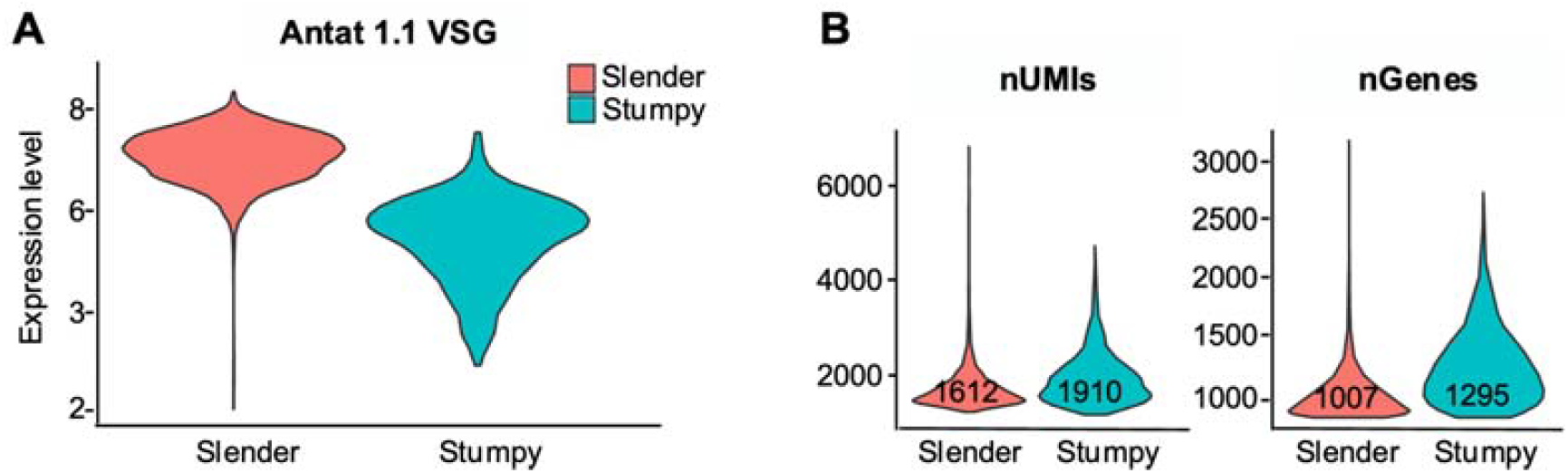
Stumpy-classified cells display a lower VSG expression level, but a higher number of genes and transcripts. **(A)** Violin Plot of expression level of the active VSG AnTat1.1 in slender and stumpy clusters shown in Figure 2A. **(B)** Violin Plot of number of unique molecular identifiers (UMIs), and number of genes detected in each cluster shown in Figure 2A.

**Figure S2:**
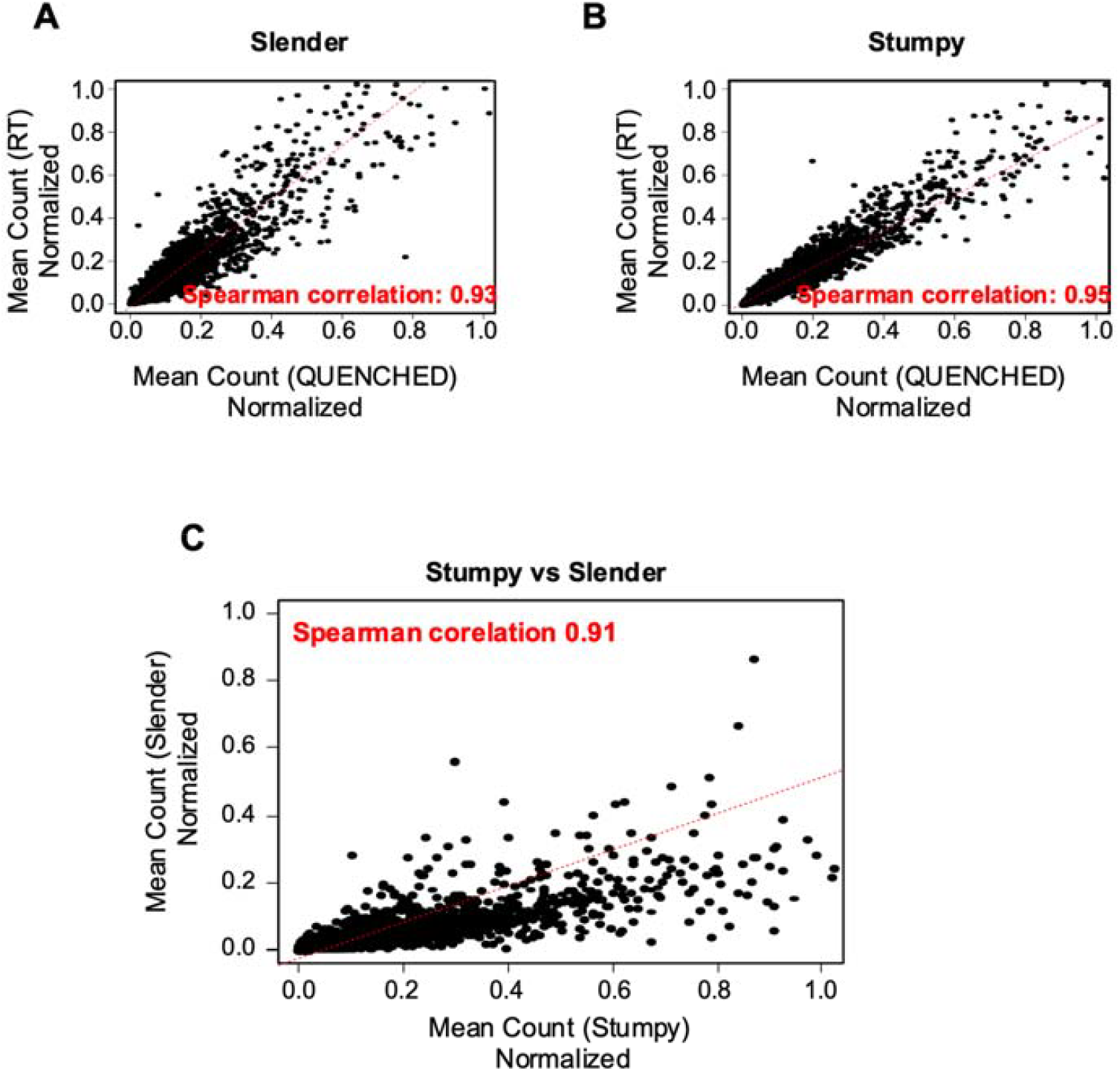
Global transcriptome comparison between slender and stumpy forms. **(A)** Spearman correlation plot of gene expression (mean gene counts-normalized) of stumpy forms in QUENCHED vs RT with a value of 0.93. **(B)** Spearman correlation plot of gene expression (mean gene counts-normalized) of slender forms in QUENCHED vs RT with a value of 0.95. **(C)** Spearman correlation plot of gene expression (mean gene counts-normalized) of stumpy vs slender with a value of 0.907.

**Figure S3:**
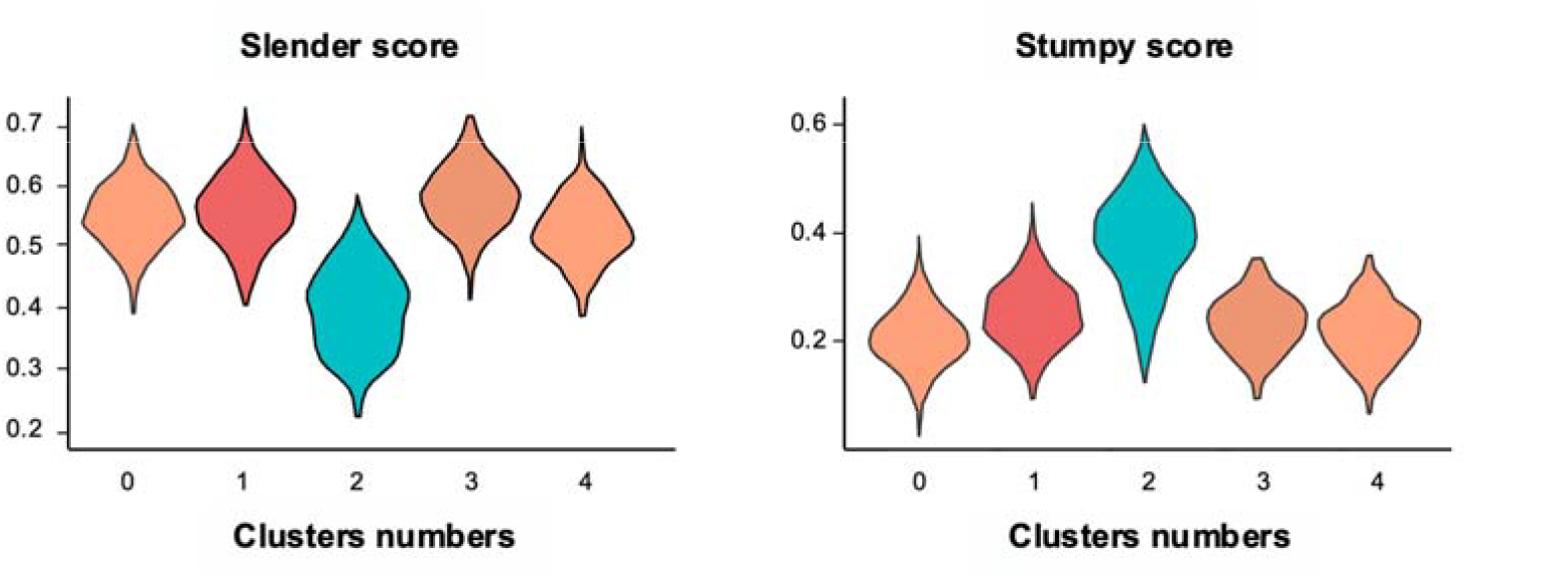
Slender and stumpy scores of in vivo T. brucei bloodstream forms. Violin plots depicting the slender and stumpy scores (as defined in Figure 2B) for the five clusters identified in single-cell data from parasites isolated from blood on day 6 post-infection (clusters also shown in Figure 4A).

## ACKNOWLEDGMENTS

We would like to thank Nicolai Siegel, Luis Graça, Ugur Sezerman, Idalio Viegas, and Tomás Gomes for their helpful discussions. We are also grateful to Michal Malecki and Lúcia Serra for their comments on the manuscript, and to Emma Briggs for providing the reference genome and informatic codes. Additionally, we appreciate the support provided by the Rodent and Bioimaging facilities.

First illustration was made using BioRender (Agreement number: LI27JG3JB5). This work was supported by the Marie Skłodowska-Curie Cell2Cell ITN grant (ref. 860675), the European Research Council (ERC) under the European Union’s Horizon 2020 research and innovation programme (771714) and La Caixa Foundation Health program (HR24-00288).

## AUTHOR CONTRIBUTIONS

LLE: Writing-original draft, designed the study, carried out all experiments and data processing, analysed the data, wrote the manuscript

LMF: acquired funding, designed the study, supervised the project, and edited the paper.

AN: carried out microscopy visualization experiments

## DATA AVAILABILITY

The raw FASTQ files from the sequencing data generated in this study have been deposited in the NCBI Gene Expression Omnibus (GEO) database under accession number “SUB15094190”. Additional data related to the sequencing analysis can be found in the Supplementary Data files. All code used for the analysis is available on

GitHub: https://github.com/Laritabonita/Encapsulation.

## DECLARATION OF GENERATIVE AI AND AI-ASSISTED TECHNOLOGIES IN THE WRITING PROCESS

During the preparation of this work the author(s) used ChatGPT 4o in order to improve the readability of the text originally written by the team. After using this tool, the author(s) reviewed and edited the content as needed and take(s) full responsibility for the content of the publication.

